# An extended reconstruction of human gut microbiota metabolism for personalized nutrition

**DOI:** 10.1101/2020.10.23.350462

**Authors:** Telmo Blasco, Sergio Pérez-Burillo, Francesco Balzerani, Alberto Lerma-Aguilera, Daniel Hinojosa-Nogueira, Silvia Pastoriza, María José Gosalbes, Nuria Jiménez-Hernández, M. Pilar Francino, José Ángel Rufián-Henares, Iñigo Apaolaza, Francisco J. Planes

## Abstract

Understanding how diet and gut microbiota interact in the context of human health is a key question in personalized nutrition. Genome-scale metabolic networks and constraint-based modeling approaches are promising to systematically address this complex question. However, when applied to nutritional questions, a major issue in existing reconstructions is the lack of information about degradation pathways of relevant nutrients in the diet that are metabolized by the gut microbiota. Here, we present AGREDA, an extended reconstruction of the human gut microbiota metabolism for personalized nutrition. AGREDA includes the degradation pathways of 231 nutrients present in the human diet and allows us to more comprehensively simulate the interplay between food and gut microbiota. We show that AGREDA is more accurate than existing reconstructions in predicting output metabolites of the gut microbiota. Finally, using AGREDA, we established relevant metabolic differences among clinical subgroups of Spanish children: lean, obese, allergic to foods and celiac.

## INTRODUCTION

Understanding how diet and gut microbiota interact in the context of human health is a key question in personalized nutrition^1^. Nutrients derived from the diet affect the abundance of different species present in the gut microbiome, which, on the other hand, release key metabolites and signals that regulate host health. The relevance of this interaction is supported by an increasing body of literature showing that the beneficial effect of dietary interventions in different clinical conditions is associated with specific signatures of the gut microbiota^1–3^.

Given the complex molecular events implied in this relevant question, the development of computational models, driven by meta-omics data, constitutes a major task in Systems Biology^4,5^. In particular, the integration and analysis of genome-scale metabolic models of different bacterial species that are present in the human gut microbiota have received much attention^6^. Thanks to the tremendous effort in the last years to generate high-quality computational platforms for metabolic reconstruction^7–10^, extensive microbial community models of the human gut microbiome are now available. Currently, AGORA constitutes the largest effort in the literature, involving 818 species present in the human gut microbiota^11^.

These network-based community models, which integrate the metabolic capabilities of different bacterial species in the gut microbiome, can be analyzed via Constraint-Based Modeling (CBM)^12–14^. This approach is promising in personalized nutrition and could help in elucidating how different microbial species in the human gut exploit and transform nutrients derived from the diet and in systematically designing effective dietary strategies when the gut microbiome is dysregulated. For example, AGORA has been already applied to predict dietary supplements for Crohn’s disease^15^. Using a similar approach, we predicted the effect of solid diet on the gut microbiota metabolism of infants^16^.

However, a major issue of current metabolic reconstruction platforms is the limited information about degradation pathways of key diet-derived nutrients. For example, AGORA only involves 99 out of 650 nutrients included in i-Diet, a commercial software for personalized nutrition^17^. In addition, universal metabolic databases, such as the Model SEED^7^, on which reconstruction platforms rely for gap filling, are incomplete and include metabolic capabilities of species that are not present in the human gut. Overall, these limitations restrict the scope of CBM approaches to establish personalized nutrition programs.

In this article, using a combination of bioinformatic tools, literature and expert knowledge, we extend AGORA and substantially improve the coverage of the metabolism of diet-derived nutrients. Particularly, we include the degradation pathways of 231 nutrients (not present in AGORA), from which 211 are phenolic compounds, a family of metabolites highly relevant for human health and nutrition. Our reconstruction, called **AGREDA** (AGORA-based REconstruction for Diet Analysis), is thus more amenable to analyze the effect of dietary interventions.

To illustrate our contribution, we first show that AGREDA outperforms AGORA in differentiating 20 typical recipes of the Mediterranean diet, according to their nutrient composition. In addition, using 16S rRNA sequencing data, we apply AGREDA to analyze the metabolic output of the gut bacterial community during the *in vitro* fermentation of lentils with faeces of children belonging to different clinical groups: lean, obese, allergic to foods and celiac. We provide experimental validation for 10 different output phenolic compounds and establish important metabolic differences in the gut microbiota of the children analyzed, emphasizing the insights derived from AGREDA that could not be obtained with AGORA. In conclusion, AGREDA addresses the necessary intersection between human nutrition, genomics and computational modeling to reach the 21^st^ century nutrition: personalized nutrition.

## RESULTS

We present a new metabolic reconstruction of the human gut microbiota that is focused on covering significant gaps in the degradation of diet-derived nutrients. We started from AGORA^11^, the most detailed metabolic resource that includes 818 reconstructions of bacterial species present in the human gut microbiota. We first assessed the number of nutrients present in i-Diet^17^, a commercial nutritional software designed to elaborate optimal diets, that are annotated in AGORA. We found that 551 out of 650 metabolites are missing, which justifies the need for the work presented here.

In order to reduce the size of the reconstruction and computation time, we did not take into account the boundaries of individual organisms and extracted a non-redundant set of reactions and metabolites from AGORA. In other words, we defined a metabolic network with only two compartments: external and internal, obtaining 2473 metabolites and 5312 reactions. Note here that, for meta-omics data integration, we stored for each reaction its taxonomic annotation in AGORA. Henceforth, this summarized network is referred to as AGORA.

We then built a universal metabolic network based on the Model SEED^7^ (SEED) database and expert nutritional knowledge. Through their EC numbers (if available), reactions were annotated to species present in AGORA using different bioinformatics tools and metabolic databases (Figure 1 and Methods section for details). This universal network was consistently integrated with the reactions and metabolites from AGORA. We finally applied a gap filling algorithm to include in our reconstruction the maximum number of diet-derived nutrients and their degradation pathways, which was based on FastCoreWeighted, included in the COBRA Toolbox^18,19^ (see Methods section for full details). Our final reconstruction is called AGREDA (AGora-based REconstruction for Diet Analysis, Supplementary Data 1).

**Figure 1.**
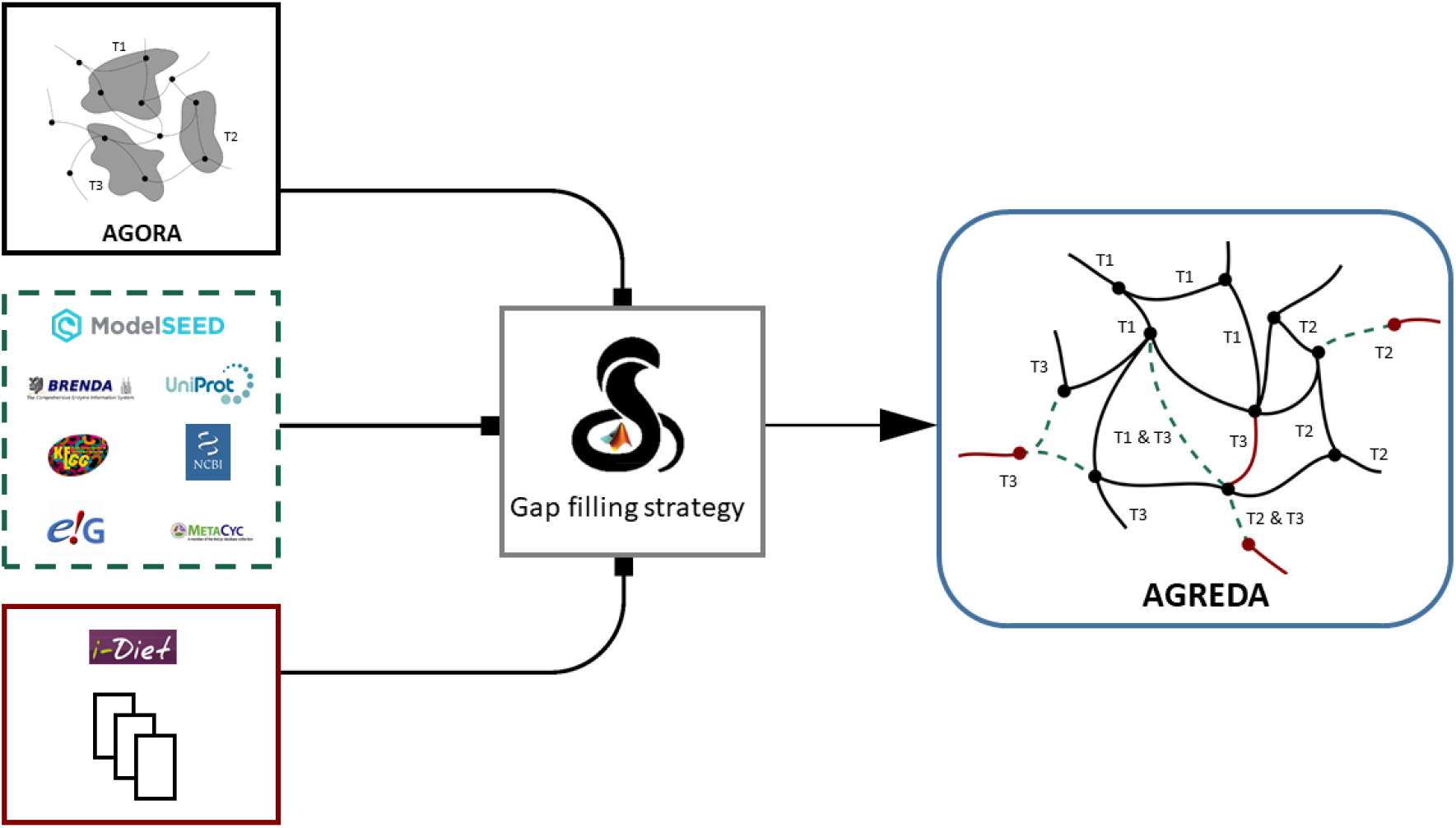
Summary of the reconstruction pipeline. First, AGORA reconstructions^11^ (black) are combined into a supra-organism model, while saving the taxonomic assignment of the reactions. Next, Model SEED^7^ reactions (green) are annotated to AGORA species through EC number information (see Methods section) and added to the supra-organism model. Then, metabolites provided by i-Diet and manually curated by expert nutritional knowledge are integrated with AGORA and Model SEED (maroon). Finally, gap-filling techniques, based on the Cobra Toolbox^18,19^, are applied to derived AGREDA.

AGREDA adds to AGORA 899 reactions and 401 metabolites, from which 231 are diet-derived nutrients from i-Diet not included in AGORA. Full details, including functional and taxonomic annotation of reactions, can be found in Supplementary Data 2. Figure 2a shows the number of reactions and metabolites related to each species grouped by the respective phyla. It can be observed that all phyla contain a higher number of metabolites in AGREDA than in AGORA. Specifically, each phylum in AGREDA contains on average 70 reactions and 170 metabolites more than in AGORA.

**Figure 2.**
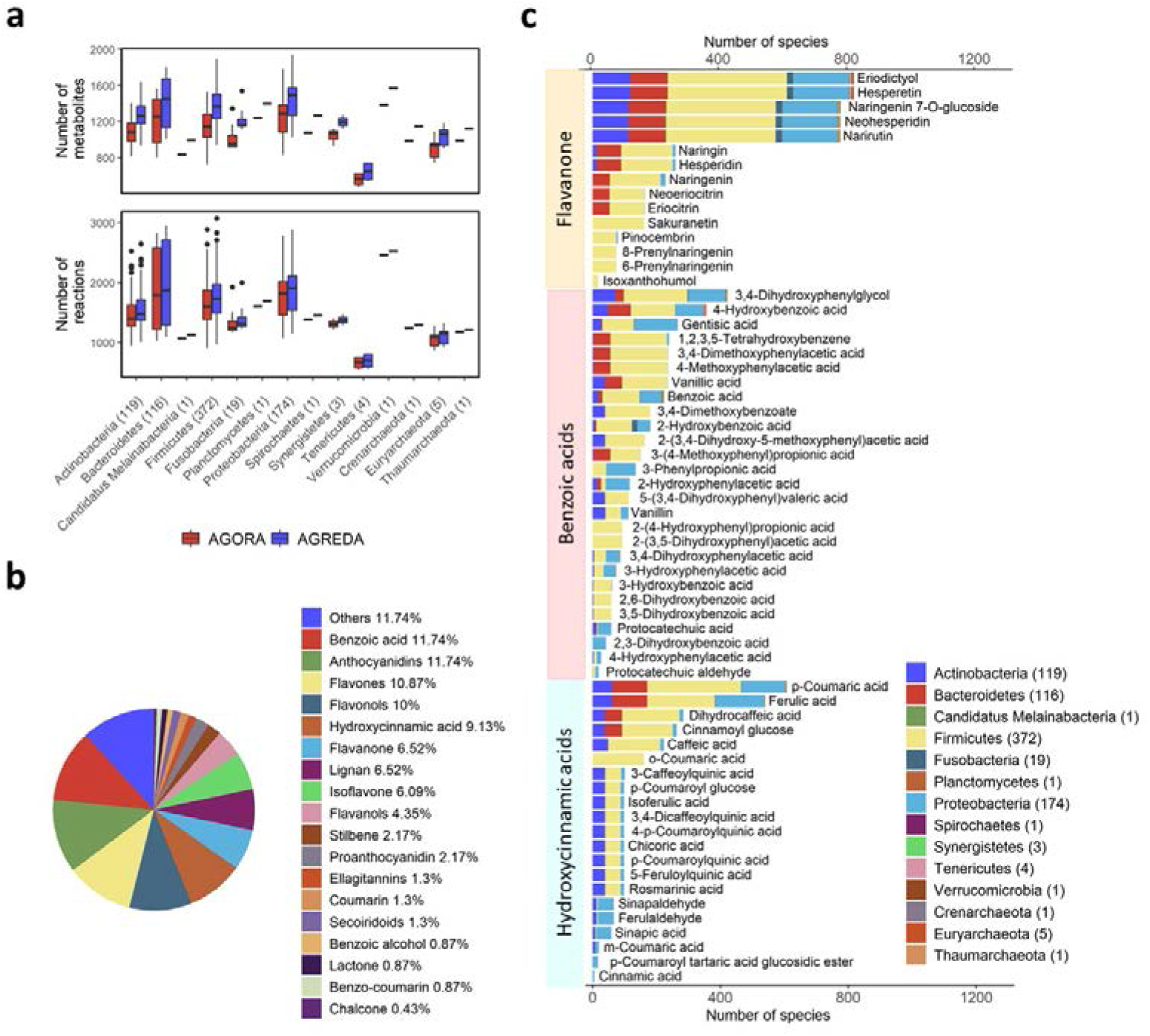
Main features of AGREDA. **(a)** Number of metabolites and reactions in AGORA (red) and AGREDA (blue) belonging to the 14 phyla present in the models. The number of strains per phylum is shown in brackets. **(b)** Distribution of the 211 phenolic compounds added by AGREDA separated in 19 families. **(c)** Degradation capabilities for three families of phenolic compounds present in AGREDA. The total number of strains in each phylum is reported in brackets.

An important set of metabolites included in AGREDA is that of phenolic compounds. These nutrients are widespread in the vegetal kingdom, where they act as a defensive system against external aggressions and have been pointed out to be responsible for many of the health benefits of vegetable consumption. AGREDA covers a very wide range of phenolic compounds, from the simpler ones (benzoic and hydroxycinnamic acids) to the more complex (proanthocyanidins), with all families represented (Figure 2b). Overall, AGREDA added the degradation pathways of 211 phenolic compounds, significantly improving the coverage of AGORA, which only contained 19 phenolic compounds.

The daily intake of phenolic compounds is rather high, since they are especially abundant in highly consumed food items such as tea or coffee (specially rich in cinnamic acids and flavan-3-ols) and fruits, vegetables and legumes (wide range of different flavonoids)^20^. However, they are barely absorbed in the small intestine and reach the gut microbiota where they are metabolized by organisms belonging to different phyla, usually into smaller molecules that are more easily absorbed in the large intestine^21^. Therefore, the benefits of most phenolic compounds are actually exerted by their output metabolites, hence the importance of being able to define their microbial metabolization^22^. Figure 2c, for example, shows the degradation capabilities of different phyla for 3 families of phenolic compounds: flavanones, benzoic acids and hydroxycinnamic acids.

In order to assess the improvement over AGORA, we selected 20 representative recipes and employed i-Diet to calculate the nutrients present in each of them (Supplementary Data 3). As shown in Figure 3a, approximately only half of the nutrients of each recipe captured by AGREDA are captured by AGORA. In addition, the heatmaps in Figure 3b represent the dissimilarity (Jaccard’s distance) among the sets of nutrients present in each recipe captured by AGORA and AGREDA, respectively. We observe that the latter is significantly greater than the former, meaning that AGREDA performs better at capturing the potential metabolic differences between the recipes. We can, therefore, conclude that AGREDA provides us with a more accurate tool to assess the effects of the different diets on the gut metabolism with a straightforward application to personalized nutrition.

**Figure 3.**
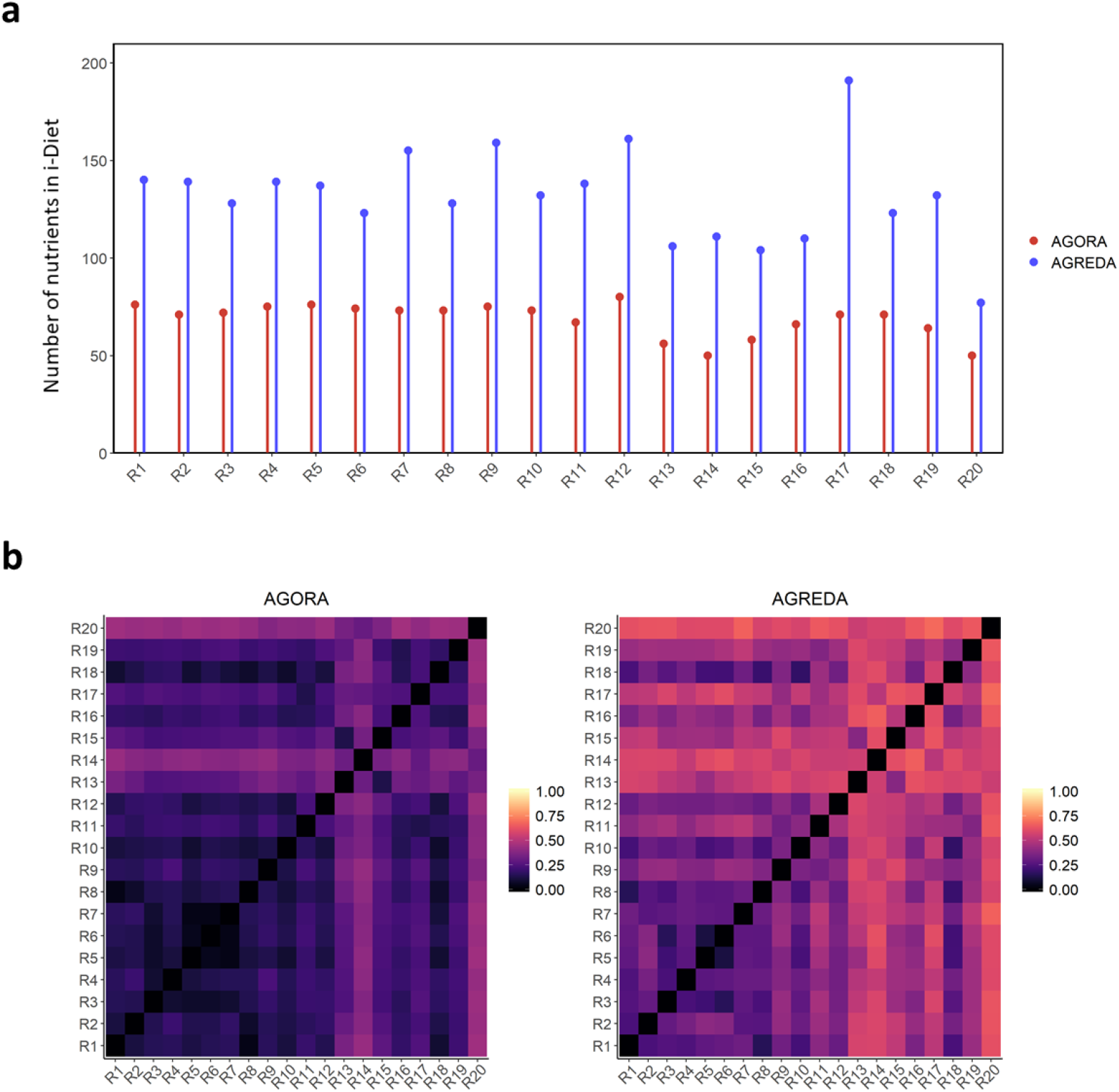
Capability of the models to capture the nutritional composition of 20 representative recipes. **(a)** The number of nutrients that AGORA and AGREDA are able to capture per recipe. Note that all the metabolites present in AGORA are also included in AGREDA. **(b)** Differences between the nutritional content of the recipes captured by AGORA and AGREDA respectively. The Jaccard’s distance between the composition of the recipes is represented.

### *In vitro* fermentation of lentils with children faeces

A commercial Spanish recipe of boiled lentils was used for the next set of experiments. Its nutritional composition was obtained by means of the i-Diet software^17^ (Supplementary Data 3). Lentils were fermented *in vitro* with faecal inocula from children belonging to 4 different clinical conditions, *i.e*. lean, obese, allergic to foods and celiac. Seven inocula were prepared with the fecal samples proceeding from lean, obese and celiac children, while six were prepared with those proceeding from allergic children, for a total of 27 fermentations. The taxonomic composition of the microbiota present in the different fermentations was measured by means of 16S sequencing technologies (see Methods section). Next, we contextualized the reference AGREDA and AGORA models with the nutritional information of the lentils recipe and the taxonomic composition of the fecal inocula (see Methods section, Supplementary Data 3), obtaining 27 context-specific AGORA and 27 context-specific AGREDA models for the aforementioned conditions.

We experimentally measured the presence or absence of a set of 10 representative phenolic compounds in three inocula per clinical condition through targeted metabolomics analysis, for a total of 12 samples (see Methods section for details, Supplementary Data 3) and assessed the predictive potential of the respective AGORA and AGREDA models (Figure 4a). Here, we noticed that the reference (uncontextualized) AGORA network only captures 3 out of 10 measured phenolic compounds, while the reference (uncontextualized) AGREDA network contains all the measured metabolites. As a consequence, the sensitivity of the AGREDA context-specific models is remarkably higher than that of the AGORA context-specific models (76,4% versus 22,3%, Figure 4a). Moreover, AGREDA outperforms AGORA regarding accuracy (75% versus 32,5%). We, therefore, conclude that the new metabolites and degradation pathways included in AGREDA significantly improve our predictive capacity of gut microbiota metabolism and enable the detection of output metabolites not considered in AGORA.

**Figure 4.**
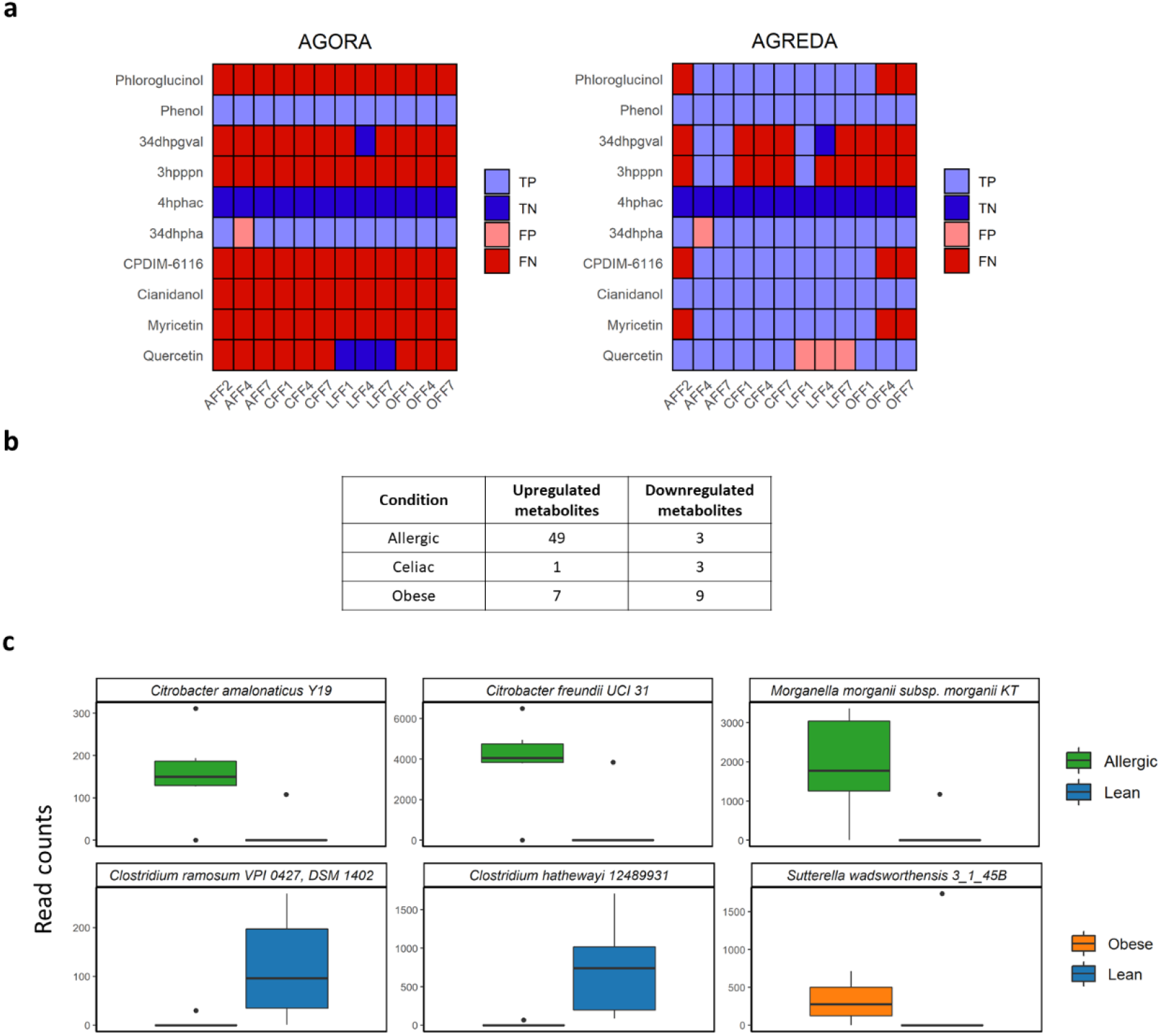
Case Study: Degradation of a traditional lentils recipe. **(a)** Comparison of the predictive potential of 10 metabolites secreted by the gut microbiota between AGREDA and AGORA. The medium was defined by the nutrients of a traditional lentils recipe. **(b)** Summary of the AGREDA prediction of representative metabolites secreted by the gut microbiota in samples from allergic, obese, celiac children in comparison to lean children. **(c)** Bacterial species involved in the biosynthesis of histamine (green), tryptamine (green), myricetin (blue) and isoprene (orange). Rarefaction was applied for normalization. Abbreviations: TP (True Positives), TN (True Negatives), FP (False Positives), FN (False Negatives), 34dhpgval (5-(3’,4’-Dihydroxyphenyl)-gamma-valerolactone), 3hpppn (3-(3-hydroxy-phenyl)propionate), 4hphac (4-hydroxyphenylacetate), 34dhpha ((3,4-dihydroxyphenyl)acetate), CPDIM-6116 (Dihydrocaffeic acid).

Next, we employed the aforementioned AGREDA contextualized reconstructions aiming at identifying the relevant output metabolites for each disease condition in comparison to the lean state (Bayesian Logistic Model, p-value <= 0.05) using the 27 samples previously described. We found a list of 52, 16 and 4 relevant metabolites for allergic, obese and celiac conditions, respectively (Figure 4b, Supplementary Data 3). Importantly, 22 out of these 72 metabolites were only captured in AGREDA and not in AGORA.

The samples from allergic children show more extreme differences with respect to the rest of conditions. In particular, we identified the biosynthesis of histamine and tryptamine as a specific feature in these fermentation samples. Both metabolites are closely related since tryptamine tends to increase the levels of histamine in the organism^23^ and the latter is involved in the inflammatory response of allergies^24,25^. Interestingly, the gut microbiota supports the production of histamine and tryptamine, as reported in different works for adults^26,27^. We could identify the species involved in histamine and tryptamine biosynthesis, namely, *Citrobacter amalonaticus Y19, Citrobacter freundii UCI 31* and *Morganella morganii subsp. morganii KT*, which are only present in these samples (Figure 4c).

Regarding the obese children’s samples, two relevant metabolites caught our attention, namely, isoprene and myricetin. The former is predicted to be produced exclusively in the obese condition and, interestingly, it has been reported to be involved in many metabolic disorders and as a potential obesity marker exhaled in breath^28,29^. With respect to myricetin, however, just the opposite occurs since it is only produced in the fermentations with inocula from the lean children’s stools but not in those from the obese children. Importantly, several works performed with mice have shown that this phenolic compound provides anti-obesity effects^30,31^. Note that myricetin is one of the metabolites that would not have been captured by AGORA. For both isoprene and myricetin, we could identify the species involved in their biosynthesis, *i.e. Sutterella wadsworthensis 3_1_45B* and *Clostridium hathewayi 12489931* and *Clostridium ramosum VPI 0427, DSM 1402*, respectively (Figure 4c).

## DISCUSSION

Constraint-based modeling constitutes a promising approach to investigate the interaction of diet and gut microbiota and their impact in the host’s health. In the last years, the number of high-quality genome-scale metabolic reconstructions of species present in the human gut has significantly increased, aiming to conduct a more comprehensive analysis of the gut microbiota metabolism. However, they need further developments to become a practical tool in the area of personalized nutrition, since a large variety of key nutrients present in the diet are not considered in these reconstructions. This limitation could substantially impair our study of the interplay between diet and gut microbiota metabolism.

In this article, we directly address this relevant issue and extend AGORA^11^, the largest repository of metabolic reconstructions of species present in the human gut microbiome. In particular, we add to AGORA the degradation pathways of 231 nutrients included in i-Diet^17^, a commercial nutritional software designed to elaborate optimal diets, collectively involving 899 new reactions and 401 new metabolites. Our reconstruction, termed AGREDA, was built through an exhaustive literature analysis and gap filling algorithms using the Model SEED^7^ as universal database. For this task, we used different bioinformatic tools to integrate SEED and AGORA and avoided the use of reactions with limited evidence in the human gut microbiota. As a result, our proposed reactions in AGREDA include taxonomic annotation to species present in AGORA, which facilitates the analysis of their activity with 16S rRNA sequencing data.

Note here that we decided to follow a supra-organism strategy to build AGREDA. This was done to reduce the size of the community model and, therefore, the computation time of our simulations. Given our positive results, this simplification does not seem to affect our predictions. However, a future study should analyze the deviations derived from our supra-organism assumption and, if necessary, correct AGREDA to include exchange reactions and boundaries among the different species involved.

AGREDA focuses on phenolic compounds. This family of compounds is one of the most abundant source of bioactive compounds present in the human diet, mainly in plant foods, fruits and plant-derived beverages, which are mostly metabolized by the gut microbiome. With the inclusion of the degradation pathways of more than 200 nutrients, AGREDA constitutes the largest effort in the literature to compile the metabolism of phenolic compounds in the human gut microbiome. Despite our advance, there is substantial room for improvement, since AGREDA currently only includes 99 out of 372 metabolites detailed in Phenol-Explorer, the first comprehensive database of polyphenol contents in foods. Many of them are not annotated in universal metabolic databases, such as KEGG^32,33^ or SEED, requiring new strategies to address this issue. In this direction, enzyme promiscuity methods constitute a promising approach to further complete degradation pathways of phenolic compounds.

Importantly, AGREDA more accurately models the effect of diet on gut microbiota metabolism than AGORA, as shown in Figure 3 for 20 different representative recipes, where a significantly better coverage of their nutrient composition was obtained. This advance logically allows us to carry out a more comprehensive analysis of output metabolites from the gut microbiota. This was illustrated in the case study of lentils, where AGREDA showed higher accuracy than AGORA in predicting 10 experimentally measured output metabolites.

Finally, we applied AGREDA to assess metabolic differences in the way the gut microbiome of different clinical groups of children degrade lentils. We identified relevant insights for allergic and obese children when compared with the lean condition, but limited evidence for differences in the metabolic output of the microbiota of the celiac children. We found supporting literature for some of our predictions, particularly histamine and tryptamine for fermentations with inocula from the allergic children, and isoprene and myricetin for the obese children. Further experimental validation is necessary to confirm our predictions in a larger cohort of children. However, our work opens new avenues to incorporate the effect of gut microbiota in personalized nutrition programs.

## METHODS

### Universal biochemical reaction database

We start from AGORA^11^, which comprises manually curated metabolic models of 818 species of the human gut microbiome. In order to reduce the computational cost, we followed a supra-organism strategy and removed the boundaries between different species. Based on AGORA, we defined a non-redundant set of metabolites and reactions, including their taxonomic assignation. Overall, we obtained 2473 metabolites and 5312 reactions.

AGORA currently lacks the degradation pathways of key diet-derived metabolites. In particular, we found that AGORA only includes 99 out of 650 diet-derived metabolites from i-Diet^17^, a commercial nutritional software designed to elaborate optimal diets. Among these neglected metabolites, we found an important number of phenolic compounds, whose functional role in the human gut microbiota is of major interest in the field of personalized nutrition^34^. To overcome this issue, we integrated the information provided by AGORA with the Model SEED database^7^ (SEED), as well as with other metabolic databases and expert knowledge of gut microbiota metabolism, as we detail below.

We first downloaded SEED from the online portal (https://modelseed.org/), which involves 20133 metabolites and 34655 reactions. To minimize the inclusion of reactions from species not active in the human gut microbiota, we decided to annotate the EC numbers present in SEED with the species present in AGORA. Note here that SEED does not incorporate Gene-Protein-Reaction rules, as available in AGORA; instead, SEED presents a wide functional annotation of reactions through EC numbers. In this event, the integration of SEED into AGORA can be done through the taxonomic annotation of its EC numbers. We describe below the different strategies followed to carry out this task with existing genomic annotation tools and relevant metabolic databases.

Genome *fasta* files from different species in AGORA were downloaded from GenBank^35^ and Ensembl^36^ through the NCBI taxonomy identifier and species name, respectively. These genomes were annotated using myRAST software from the RAST Server^37^, which outputs their protein-encoding genes and (if available) associated EC numbers. This information was incorporated into the reactions present in SEED. In addition, from the KEGG database^32,33^, we downloaded the list of EC numbers for 500 species present in AGORA. With this information, we could further annotate reactions in SEED without taxonomic information.

We also performed a manual annotation of reactions and EC numbers present in SEED. We found that several reactions that did not contain any EC number information in SEED were annotated in public databases such as KEGG or MetaCyc^38^. Based on them, we extracted more reactions with enzymatic information and repeated the process described above for taxonomic annotation. For the remaining EC numbers without taxonomic information, we manually looked for additional information in KEGG, BRENDA^39^ and UniprotKB^40^ databases. After this process, we obtained a list of 3577 different EC numbers and 14021 reactions in SEED that are related to at least one of the species in AGORA.

We noticed that some metabolites in SEED were involved in reactions under different names. Using both manual curation and chemoinformatic tools, we identified and deleted metabolites and reactions that were duplicated in SEED. In particular, we first extracted the InChI identifier for the metabolites in SEED (13028 out of 20133 metabolites), based on PubChem^41^, the Human Metabolome Database^42^, KEGG and RetroRules database^43^. We then conducted a similarity analysis with RDKit package^44^ and the Morgan (circular) fingerprint with radius 2^45^. Fingerprints with similarity 1 were obtained and manually checked. We removed 703 repeated metabolites and 1054 reactions from SEED.

In order to integrate AGORA and SEED, we performed an automatic search of the compound names in both sources and identified duplicated metabolites and reactions. SEED added to AGORA 17820 metabolites and 32409 reactions, including 12459 with taxonomic assignment.

In addition, we manually identified the list of nutrients from i-Diet present in SEED, finding 232 that were not present in AGORA. We created an exchange reaction for each of these nutrients and included them in our metabolic database. We also added 221 reactions and 19 metabolites from expert knowledge and existing literature of metabolism of phenolic compounds in the gut microbiota, including their taxonomic annotation (Supplementary Data 2). After this final step, our universal biochemical reaction database reached 20376 metabolites and 38059 reactions. Note here that 20023 and 13478 of these reactions do not have taxonomic and functional assignment, respectively.

### Gap filling strategy

Our aim is to extend AGORA and include the missing metabolic pathways of the 232 diet-derived nutrients using the least possible information from our universal biochemical database described above. Note here that 211 out of these 232 nutrients are phenolic compounds, which are particularly interesting in personalized nutrition.

In order to fill these gaps, we used the implementation of FastCoreWeighted included in the COBRA Toolbox^18,19^. This reconstruction algorithm requires the definition of a subset of reactions that must take part in the resulting network, termed core, and efficiently identifies the reactions needed from the universal database to make the core functional. In addition, it allows us to penalize differently the inclusion of reactions from our universal database. Here, we set a weight equal to 0 for reactions in the core, 0.1 for reactions with taxonomic assignment to species in AGORA, 50 for reactions without taxonomic assignment but with functional annotation (at least one EC number available), 100 for reactions without taxonomic and functional annotation, and 1000 for reactions manually assigned to plant metabolism.

As we found dependencies between different nutrients from i-Diet, namely some of them are interconnected as inputs and outputs, we run FastCoreWeighted sequentially, updating the core at each iteration. In the first iteration (Iteration 1), the core included the reactions from expert knowledge and AGORA. In the second iteration, the core comprised the resulting network from Iteration 1 and the input exchange associated with the first nutrient from i-Diet. In the third iteration, the core comprised the resulting network from Iteration 1 and the input exchange associated with the second nutrient from i-Diet. This process was repeated for the 232 nutrients from i-Diet. Reactions obtained at each iteration were included in the final model.

Note here that, in order to include each input exchange reaction as part of the core in the different iterations, we split reversible reactions in our universal database into two irreversible steps. In addition, when we added the input exchanges of nutrients from i-Diet in the different iterations described above, we penalized the inclusion of their associated output exchanges to avoid artifacts in the resulting network (weight of 1e5). The same approach was employed for the output exchanges of i-Diet metabolites.

We integrated the reactions selected in the different iterations described above, obtaining an active network made of 2920 metabolites and 6277 reactions. At this stage, we still had 51 reactions without taxonomic assignment. To avoid false positives, we deleted this subset of reactions and ran fastFVA^46^, obtaining a metabolic model, called AGREDA (AGORA-based reconstruction for diet analysis), that involves 2744 metabolites and 6112 reactions. AGREDA can degrade and produce 207 and 208 (out of 232) metabolites from i-Diet, respectively. Full details can be found in Supplementary Data 2.

### Contextualization of AGORA and AGREDA for different clinical conditions

In order to obtain the context-specific models for the given conditions, the same methodology was applied to both AGORA and AGREDA. First, the uptake of those nutrients that were not present in the recipe was blocked, by setting the lower bound of the respective reactions equal to zero. Next, by means of the 16S sequencing data, all those reactions which were not related to at least one taxon present in the given sample were blocked by setting both their lower and upper bounds equal to zero. Finally, fastFVA was applied and blocked reactions were removed.

### *In vitro* gastrointestinal digestion and fecal fermentation of lentils

For the *in vitro* digestion and fermentation, the following reagents were used: potassium di-hydrogen phosphate, potassium chloride, magnesium chloride hexahydrate, sodium chloride, calcium chloride dihydrate, sodium mono-hydrogen carbonate, ammonium carbonate, hydrochloric acid, all obtained from Sigma-Aldrich (Germany). The enzymes – salivary alpha-amylase, pepsin from porcine, and bile acids (bile extract porcine) – were purchased from Sigma-Aldrich, and porcine pancreatin was from Alfa Aesar (United Kingdom). The fermentation reagents (sodium di-hydrogen phosphate, sodium sulfide, tryptone, cysteine, and resazurin) were obtained from Sigma-Aldrich (Germany).

The *in vitro* digestion method was carried out according to the protocol described by Brodkorb and colleagues^47^. Briefly, in the oral phase, 5 mL of salivary solution with alpha-amylase (75 U/mL) and 25 μL of 0.3 M CaCl_2_ were added to 5 g of lentils and the mix was incubated at 37°C for 2 minutes. Then, 10 mL of gastric solution with pepsin (2000 U/mL) and 5 μL of 0.3 M CaCl_2_ were added and the pH was lowered to 3.0 by adding 1N HCl; the mix was then incubated at 37°C for 2 hours. Finally, 20 mL of intestinal solution with pancreatin (100 U/mL), bile salts (10 mM) and 40 μL of 0.3 M CaCl_2_ were added and the pH was raised to 7.0 with 1N NaOH, after which the mix was incubated at 37°C for 2 hours. The enzymatic reactions were halted by immersing the tubes in iced water. The samples were then centrifuged at 6000 rpm for 10 minutes at 4°C and the supernatants separated from the solid residue or pellet.

The *in vitro* fermentation was carried out according also to the protocol described by Pérez-Burillo et al^48^. Faeces were collected from three children (9-11 years old) from each of the groups studied: cow’s milk allergic, celiac, obese (BMI >= 30) and lean (BMI <= 25). Faeces from children belonging to the same group were pooled together to reduce inter-individual variability. Additionally, seven different inocula were prepared from the celiac, lean and obese derived pools respectively and six different inocula from the allergic derived one, yielding therefore a total of 27 fermentation experiments. Right after collection, faeces were mixed with glycerol (50:50 w/v) and frozen at −80°C. Briefly, 500 mg of digested wet-solid residue were placed in a screw-cap tube. The 10% of the digestion supernatant was added to the solid residue in order to mimic the fraction that is not readily absorbed after digestion. Then, 7.5 mL of fermentation medium (15 g/L of peptone, 0.312 mg/L of cysteine and 0.312 mg/L of Na2S, adjusted to pH 7.0) and 2 mL of inoculum (consisting of a solution of 32% faeces in phosphate buffer 100 mM, pH 0 7.0) were added, to reach a final volume of 10 mL + digestion supernatant volume. Nitrogen was bubbled through the mix to produce an anaerobic atmosphere and the mix was then incubated at 37°C for 20 hours under oscillation. Immediately afterwards, the samples were immersed in ice, to stop microbial activity, and centrifuged at 6000 rpm for ten minutes. The supernatant was collected as a soluble fraction potentially absorbed after fermentation and stored at −80°C.

### DNA extraction and amplicon sequencing

Genomic DNA from the solid residues of the fermentation reactions was extracted using the MagNaPure LC JE379 platform (ROCHE) and the DNA Isolation Kit III (Bacteria, Fungi) Ref 03264785001, following manufacturer’s instructions, with a previous lysozyme lysis. DNA quality was determined by agarose gel electrophoresis (0.8 % wt/vol agarose in Tris-acetate-EDTA buffer) and quantified using the Qubit 3.0 Fluorometer (Invitrogen) and the Qubit dsDNA HS Assay Kit.

In order to prepare amplicon libraries, DNA at 5ng/μL in Tris 10mM (pH 8.5) was used for the Illumina protocol for the small subunit ribosomal DNA gene (16S rRNA) Metagenomic Sequencing Library Preparation (Cod 15044223 Rev. A). PCR primers targeting the V3-V4 hypervariable region of the 16S rRNA gene were designed as described by Klindworth and colleagues^49^, i. e. forward primer (5’-TCGT CGGC AGCG TCAG ATGT GTAT AAGA GACA GCCT ACGG GNGG CWGCA-G3’) and reverse primer (5’-GTCT CGTG GGCT CGGA GATG TGTA TAAG AGAC AGGA CTAC HVGG GTAT CTAA TCC3’). Primers were fitted with adapter sequences added to the gene-specific sequences to make them compatible with the Illumina Nextera XT Index Kit (FC-131-1096). After 16S rRNA gene amplification, amplicons were multiplexed and sequenced in an Illumina MiSeq sequencer according to the manufacturer’s instructions in a 2 x 300 cycles paired-end run (MiSeq Reagent kit v3MS-102-3001).

### Taxonomic assignment of 16S rRNA sequencing data

16S rRNA gene raw sequence reads were processed, trimmed and clustered into amplicon sequence variants (ASVs) using DADA2^50^. Once we obtained the ASV table, we assigned species-level taxonomic identifications to each ASV with DADA2, based on exact matching (100% identity) between ASVs and the reference sequences in the Silva database (version 132)^51^.

In addition, for those ASVs that were identified with DADA2 at genus level but not at species level, we applied the MegaBLAST module from BLAST^52^. Here, we required at least 97% identity for the species level assignment; however, as MegaBLAST does not take into account the previously assigned genus level, we only considered ASVs for which MegaBLAST and DADA2 classifier method assigned the same genus. Finally, ASVs with less than 0.01% of the total number of counts were removed and rarefaction was applied up to the smallest library size across samples (52923 counts) for further analysis.

Finally, we linked each of the taxa to the species present in AGORA. As the taxonomic assignment methods typically provide information at the species level but not at the strain level, each obtained taxon could be related to different strains of AGORA, where most of the taxa are defined at strain level. In this event, our analysis was conducted at species level, which is a stricter strategy.

### Identification and quantification of phenolic compounds

For individual phenolic quantification the following standards were used: phloroglucinol, phenol, 3-(3-hydroxy-phenyl)propionate, 4-hydroxyphenylacetate, (3,4-dihydroxyphenyl)acetate, dihydrocaffeic acid, cianidanol, myricetin, and quercetin were purchased from Sigma-Aldrich (Germany). 5-(3’,4’-Dihydroxyphenyl)-gamma-valerolactone was purchased from Toronto Research Chemicals (Canada). Moreover, diethyl ether for extraction was purchased from Sigma-Aldrich (Germany).

Phenolic compounds were analyzed through UV-UHPLC as described by Perez-Burillo et al.^53^, slightly modified to adapt it to UHPLC. In brief, one mL of fermentation supernatant was mixed with 1 mL of diethyl ether and kept in the dark at 4°C for 24 hours. The organic phase was then collected and another two extractions with diethyl ether were performed. These 3 mL of diethyl ether were dried in a rotary evaporator set at 30°C and the solid residue was resuspended in 1 mL of methanol:water (50:50 v/v) mix. The mixture was then ready to be injected into UHPLC system. The UHPL is an Agilent 1290 Infinity II equipped with a quaternary pump, an autosampler kept at 5°C and a diode array detector (DAD) set at 255 nm. The column used was InfinityLab Poroshell 120 Sb-Aq 2.1 x 150 mm and 1.9 micron. The flow rate was set at 0.250 mL/min for 46 minutes. Two mobile phases were used; milli-Q water with 0.1% of formic acid (A) and acetonitrile (B) with the following gradient: 0 to 28 minute from 95% of A to 60% and from 5 to 40% of B; 28-36 minute from 60% to 0% of A and from 40 to 100% of B; 36 to 41 minute from 0% to 95% of A and from 100% to 5% of B; these last conditions are kept for 5 minutes. Identification and quantification were carried out by comparing retention times obtained from pure standards (listed in reagents section). A calibration curve for each of the compounds was performed in the range of 0.1 to 25 ppm.

## Data availability

The authors confirm that the data supporting the findings of this study are available within the article and its supplementary material.

## Acknowledgements

This work was funded by the European Union’s Horizon 2020 research and innovation programme through STANCE4HEALTH project (Grant No. 816303).

## Author contributions

M.P.F, J.A.R.-H. and F.J.P. conceived this study. T.B., F.B., I.A. and F.J.P. developed the metabolic network and performed the computational analysis. S.P.B, D.H.-N, S.P. and J.A.R.-H. carried out the in-vitro fermentations and measured the phenolic compounds. M.J.G, A.L.-A., N.J.-H. and M.P.F performed the metagenomics analysis. All authors wrote, read and approved the manuscript.

